# scDoc: Correcting Drop-out Events in Single-cell RNA-seq Data

**DOI:** 10.1101/731638

**Authors:** Di Ran, Shanshan Zhang, Nicholas Lytal, Lingling An

## Abstract

Single-cell RNA sequencing (scRNA-seq) has become an important tool to unravel cellular heterogeneity, discover new cell types, and understand cell development at single-cell resolution. However, one major challenge to scRNA-seq research is the presence of “drop-out” events, which usually is due to extremely low mRNA input or the stochastic nature of gene expression. In this paper, we present a novel Single-Cell RNA-seq Drop-Out Correction (scDoc) method, imputing drop-out events by borrowing information for the same gene from highly similar cells. scDoc is the first method that involves drop-out information to account for cell-to-cell similarity estimation, which is crucial in scRNA-seq drop-out imputation but has not been appropriately examined. We evaluated the performance of scDoc using both simulated data and real scRNA-seq studies. Results show that scDoc can impute the drop-out events more accurately and robustly; specifically, it outperforms all available imputation methods in reference to data visualization, cell subpopulation identification, and differential expression detection in scRNA-seq data.

## 1 Introduction

The analysis of transcriptome expressions via RNA-sequencing technologies has achieved great growth and success in the past two decades. “Bulk” samples, pooling thousands to millions of cells by phenotype, allow us to understand gene expression variability at a population level (e.g., tissue). Bulk cell RNA sequencing (RNA-seq) has sparked development in a various aspects of RNA-seq applications, such as differential gene expression analysis, alternative splicing analysis, variants detection and allele-specific expression, and co-expression network analysis (Wang et al., 2009; Han et al., 2015). Nonetheless, bulk cell RNA-seq is limited by averaging over a population of cells, which may obscure the interests of research such as dynamic gene expression and cellular heterogeneity (Bacher and Kendziorski, 2016). Advanced single-cell RNA-sequencing (scRNA-seq) technologies are now emerging and enable the exploration of intratissue heterogeneity (Jaitin et al., 2014; Patel et al., 2014; Buettner et al., 2015; Bjorklund et al., 2016), stochastic gene expression (Deng et al., 2014; Petropoulos et al., 2016), new cell type discovery (Poulin et al., 2016; Villani et al., 2017), cell differentiation (Tang et al., 2010; Xue et al., 2013), and cellular lineage reconstruction (Trapnell et al., 2014; Treutlein et al., 2014; Blakeley et al., 2015) at single-cell resolution. Decreasing costs and maturing technologies lead to recent fast growth and expansion of scRNA-seq studies; however, it brings us both new opportunities and challenges.

One of the greatest challenges of scRNA-seq is the so-called “drop-out” phenomenon, occurring when “a gene is observed at moderate or even high expression level in one cell but is not detected in another cell” (Kharchenko et al., 2014). Additionally, it has been pointed out that not only zero expression values but also a portion of near zero expression measurements should be considered as drop-out events (i.e., the observed/measured values are lower than their true expression) (Lin et al., 2017). Drop-out events may occur during library preparation such as extremely low mRNA input or inefficient mRNA capture and cDNA amplification, or due to biological properties such as the stochastic nature of gene expression, where the expression of a gene could switch between “on” and “off” status across different cells (Shalek et al., 2013) and therefore lead to dynamic gene expression. Drop-out events, which interface with high noise and outlier events, result in scRNA-seq data challenging to analyze (Kharchenko et al., 2014; Pierson and Yau, 2015). Several statistical tools have been recently developed to account for drop-out events of scRNA-seq data analysis in terms of normalization (e.g., scran) (Lun et al., 2016), dimensionality reduction (e.g., ZIFA, CIDR, ZINB-WaVe) (Pierson and Yau, 2015; Lin et al., 2017; Risso et al., 2018), and differential expression analysis (e.g., SCDE, MAST, SC2P) (Kharchenko et al., 2014; Finak et al., 2015; Wu et al., 2018). As an alternative, imputation methods have been recently developed to recover the signal of interest by correcting the drop-outs in such a way that they could provide the potential to re-open standard RNA-seq analytic tools to scRNA-seq data (van Dijk et al., 2017; Gong et al., 2018; Huang et al., 2018; Li and Li, 2018; Moussa and Mandoiu, 2018).

Existing scRNA-seq imputation methods can be categorized into two main approaches; one, such as MAGIC (van Dijk et al., 2017) and SAVER (Huang et al., 2018), alter all gene expression values including high non-zero values, while the others, including scImpute (Li and Li, 2018), DrImpute (Gong et al., 2018) and LSImpute (Moussa and Mandoiu, 2018), only impute drop-out events (zeros or near zero counts), and non-drop-out values are not corrected. Based on the idea of heat diffusion via Markov transition matrix, MAGIC imputes drop-out events by sharing information across (or among?) similar cells. SAVER recovers drop-out values via borrowing information from similar cells based on the assumption that the count of each gene in each cell follows a Poisson-Gamma mixture distribution. Altering all expression measurements, like both MAGIC and SAVER, could potentially alter the expression not affected by drop-out, introducing bias.

DrImpute imputes drop-out values via averaging over multiple imputations based on several runs of k-means clustering (with different initials) on different similarity matrices (e.g., Spearman and Pearson correlations) (Gong et al., 2018). scImpute first detects cell subpopulations using principal component analysis (PCA) and the k-means clustering algorithm, then identifies the drop-outs for each gene in each cell by a Gamma-Normal mixture model, and finally imputes identified drop-out events by borrowing information from similar cells via a non-negative least squares regression (Li and Li, 2018). LSImpute also imputes drop-out values by borrowing information from highly similar cells recognized, but based on Jaccard similarity and Local Sensitive Hashing (Moussa and Mandoiu, 2018). It is advantageous to impute drop-out values by adopting information from similar cells, but none of these methods yet take the drop-outs into account for finding highly similar cells. Furthermore, to impute drop-out events based on the cell-to-cell similarities could be controversial, as these similarities can be distorted by the drop-out itself. In this paper, we present a new method, scDoc, which contains three steps: (1) identifies drop-out events based on the drop-out probability computed from a novel mixture model, (2) incorporates drop-out probability into cell-to-cell similarity calculation, and (3) imputes drop-out events by borrowing information from highly similar cells.

scDoc utilizes a novel Poisson-Negative Binomial (PNB) mixture model allowing us to detect drop-out events, which could be the values dropped from a relatively low expression value to zero or near zero count. In addition, we propose a new similarity metric, Weighted Cosine Similarity (WCS), which corrects the distortion of the cell-to-cell similarity calculation due to the ignorance of drop-out events. Moreover, scDoc imputes drop-out values via weighted averaging expression measurements from highly similar cells, where the cellular similarity is used as weight. This ensures that scDoc can impute drop-out values robustly, maintaining the stochastic nature of gene expression in scRNA-seq data. The details of the workflow is summarized in Figure 1. Through simulation studies and real data analyses, scDoc is compared with two existing and peer-reviewed imputation methods (DrImpute and scImpute) in terms of data visualization, cell subpopulation clustering, and gene differential expression (DE) analysis. The results suggest that our procedure, scDoc, is precise and robust in a wide range of data scenarios.

**Figure 1.**
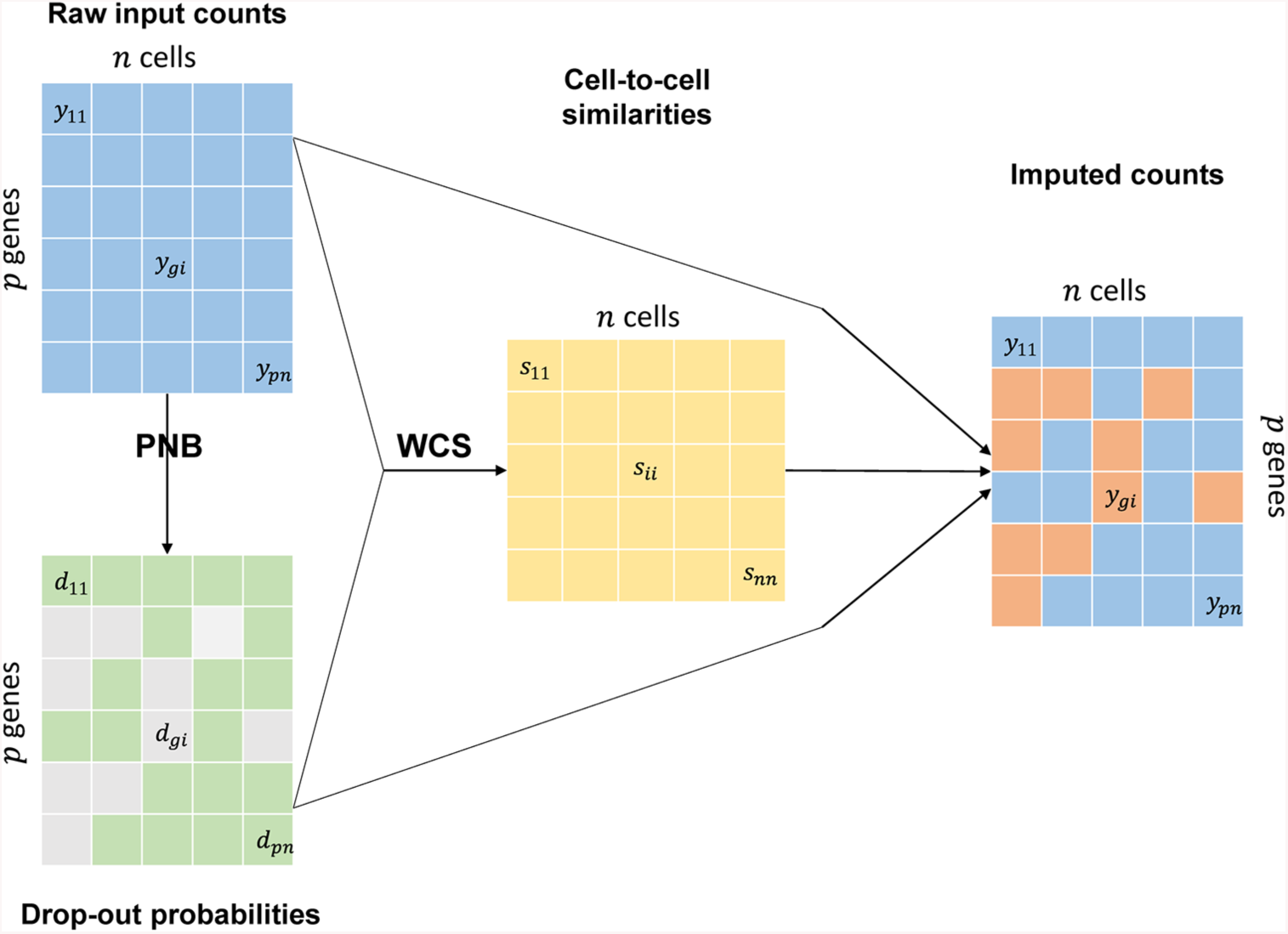
Overview of drop-outs imputation with scDoc workflow. PNB is an abbreviation for Poisson-Negative Binomial mixture distribution; WCS is an abbreviation for Weighted Cosine Similarity.

## 2 Results

### 2.1 scDoc improves the performance of visualization for scRNA-seq data

Dimensionality reduction techniques such as Principal Component Analysis (PCA) and t-Distributed Stochastic Neighbor Embedding (t-SNE) (ref) are often employed to explore and visualize high-dimensional datasets, e.g. microarray and RNA-seq data. They are naturally applied to scRNA-seq studies when scRNA-seq has emerged. However, neither of these techniques takes drop-outs into account and they were reported to lose efficiency on zero-inflated data such as scRNA-seq (Pierson and Yau, 2015; Lin et al., 2017). Recently, new statistical tools of dimensionality reduction that can address drop-out issues have been developed. For instance, ZIFA (Pierson and Yau, 2015) uses zero-inflated factor analysis to handle zero-inflated expression data, and ZINB-WaVE (Risso et al., 2018) employs zero-inflated negative binomial model to deal with both drop-out and over-dispersion. Though the performance of dimensionality reduction is improved with these methods, they only treat zeros as drop-out events and are sometimes computationally intensive. scDoc can account for drop-out event from both zero and near zero counts.

#### Results of simulation study

We simulated a series of scRNA-seq datasets. In this simulation study, decay parameter of 0.1, 0.2, and 0.3 can respectively generate around 62%, 43%, and 32% zeros into the complete data (i.e., all non-zeros) that contains three cell types. The visualization results of the first two principal components from PCA are compared for five different datasets: the complete and raw datasets, and three imputed ones via three imputation methods, scDoc, DrImpute, and scImpute. All default settings were used when applying these methods, except that the true number of cell types was given to scImpute as a prior (in order to achieve its best performance). After introducing excessive zeros, the three cell types clearly discernible in the complete dataset become ambiguous in the raw datasets and gradually indistinct with the increasing of excessive zeros (Figure. 2 second column). It was not possible to distinguish the three cell types on the 2D scatterplot when over 40% of entries were zeros in the raw datasets. All three imputation methods showed the capability to recover the distinct cell types in visualization, given about 30% excessive zeros, and scDoc outperformed the other two methods in terms of both correctness of cell type clustering and tightness of point cloud of each cell type. Though it used the true number of cell types as a prior, scImpute produced a mix of two cell types, blue and orange groups in Figure 2 (first row and fourth column). It could be because the assessment of cellular similarity in scImpute heavily depends on PCA, which is affected by even a relatively small proportion of zeros. scDoc still surpassed other methods and assisted PCA to distinguish three cell types clearly (Figure. 2, second row), though the performance deteriorated given moderate to high proportions of excessive zeros. Note that the cell types cannot be differentiated using the data imputed by scImpute when the excessive zeros were moderate. Given over 60% proportion of zeros, scDoc is the only one that can somewhat recover PCA’s ability to discriminate cell subpopulations. Data imputed with DrImpute gave no improvement compared to the raw dataset. Although there are three distinguishable clusters for the data imputed by scImpute, it was obvious that each cluster was filled with a mixture of three types of cells (Figure 2, third row). In other words, given a high proportion of excessive zeros, scImpute not only lost the power to recover true signals but also introduced bias by forcing cells into groups according to its pre-specified number of clusters. As the accuracy and efficacy of visualization deteriorate as more excessive zeros are introduced scDoc can outperform other methods due to its unique characteristics, accounting for drop-out events and eliminating their influence on cell-to-cell similarity evaluation.

Additionally, we plotted the expression profiles of these 180 genes in heatmaps for the complete and raw, and three imputed datasets with scDoc, DrImpute, and scImpute. Affirming the findings via previous 2D scatterplots, we observed that PCA’s performance of visualization worsened as more excessive zeros were introduced into the complete datasets (Appendix: Figure S1. A, B, and C). Under any scenario with different prevalence of excessive zeros, scDoc showed the capability to recover the efficacy of PCA method and dominated the performance in terms of signal recovery.

#### Real dataset 1: zebrafish data

We also used the zebrafish dataset, containing 246 single-cell transcriptomic profiling from six distinct cell types, to demonstrate scDoc’s capacity for improving performance of visualization on scRNA-seq data. The zebrafish dataset (Tang et al., 2017) contains six cells groups, among them HSPCs and HSPCs/thrombocytes come from one defined cell type but expected heterogeneity. This provides us an opportunity to separate a heterogeneous cell type into subpopulations, a common concern in real life scRNA-seq data analysis. Extremely lowly expressed genes were filtered out and 11,307 genes with least 5 counts in at least 5 cells were kept. Even though we have removed plenty of lowly expressed genes, the zebrafish dataset is still sparse with zeros composing over 87% of the total count.

Figure 3A shows the comparison of visualization performance via t-SNE on the original and three imputed datasets (we only provide t-SNE plots for this real data analysis because the main pattern cannot be visualized well in 2D PCA plots). As expected, t-SNE visualization on the log2 transformed original dataset without imputation only roughly identified clusters for neutrophils, T cells, NK cells, and B cells, whereas cells from HSCs and HSCs/thrombocytes failed to either cluster into a single defined group or separate into two distinct groups (Figure 3A). After imputation with scDoc, four undoubtedly distinct clusters can be visualized for neutrophils, T, NK, and B cells, where the group members were much more compact compared to those of the original dataset. scDoc outperformed DrImpute and scImpute in terms of cluster purity and homogeneity. For instance, t-SNE with the scDoc imputed dataset identified a pure neutrophils cluster (points in green, Figure 2B). In contrast, a few HSPCs/thrombocytes cells were blended into the neutrophils cluster when applying t-SNE to datasets imputed with DrImpute or scImpute. Though T, NK, and B cells are hypothetically similar, scImpute failed to split them clearly, while scDoc and Drimpute could differentiate them into distinct clusters. We should note that, however, DrImpute found some subpopulations for both T and NK cells, which might not be expected based on the plot of original data. The “so-called” two groups of HSPCs and HSPCs/thrombocytes cells, in the t-SNE with the scDoc imputed dataset, are loosely bundled into one single cluster. Nonetheless, they showed some heterogeneity and the data points in different groups were generally grouped towards different directions and relatively distinguishable. Contrasted with scDoc, the cell groups were scattered into several small clusters when applying t-SNE on the datasets imputed with DrImpute or scImpute. In addition, a proportion of the cells in HSPCs and HSPCs/thrombocytes types were grouped by scImpute with other cell types in a big cluster.

**Figure 2.**
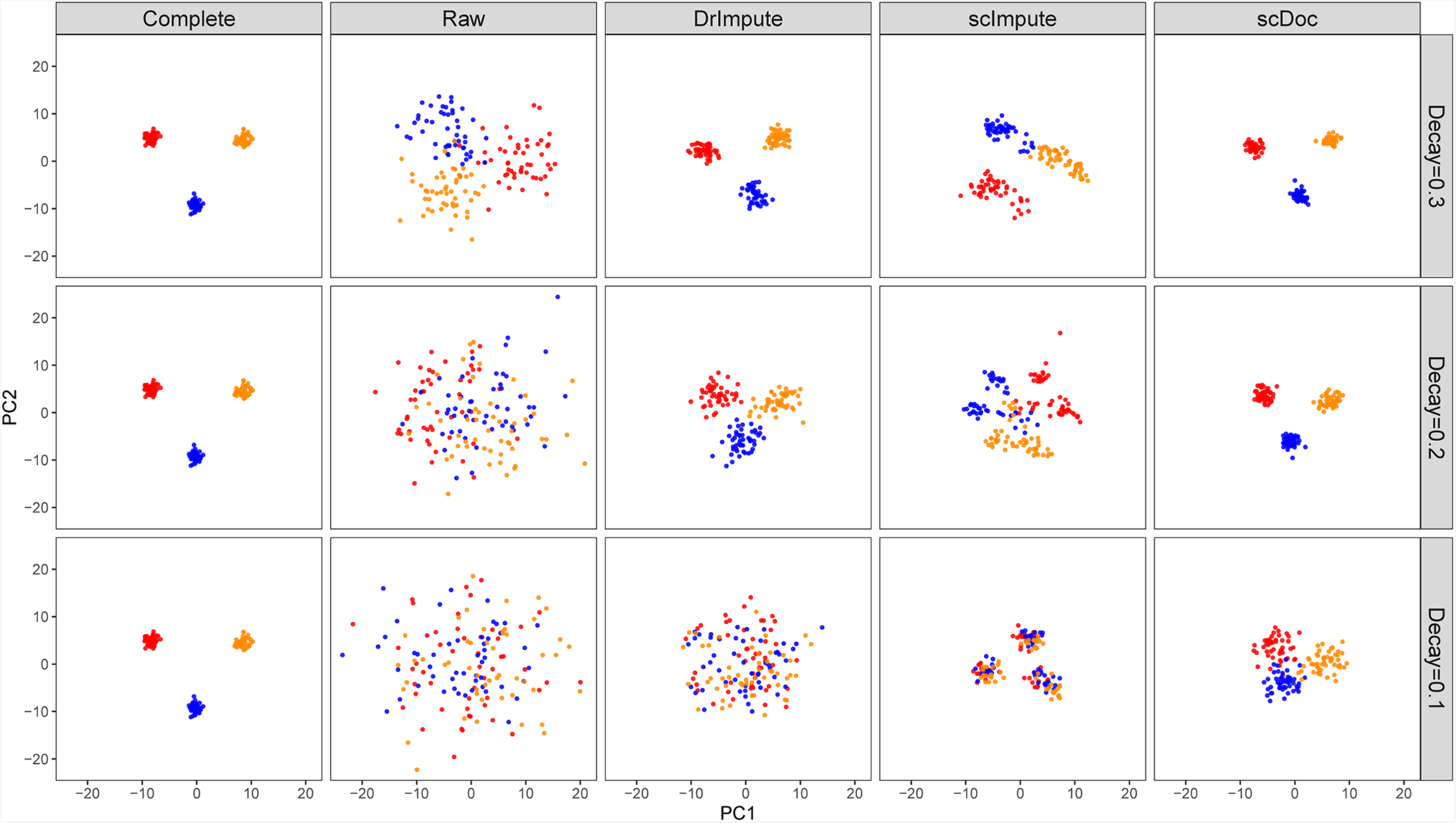
scDoc significantly improved the performance of PCA in visualizing simulated scRNA-seq count data. Plots of the first two PCs calculated from the simulated complete and raw, and three corrected datasets imputed with DrImpute, scImpute (C=3), and scDoc respectively (from left to right). Excessive zeros were introduced to the raw dataset using the double exponential function with decay parameter *ρ*. (first row) *ρ* = 0.3. The zero prevalence is ∼32%. (second row) *ρ* = 0.2. The zero prevalence is ∼43%. (third row) *ρ* = 0.1. The zero prevalence is ∼62%.

**Figure 3.**
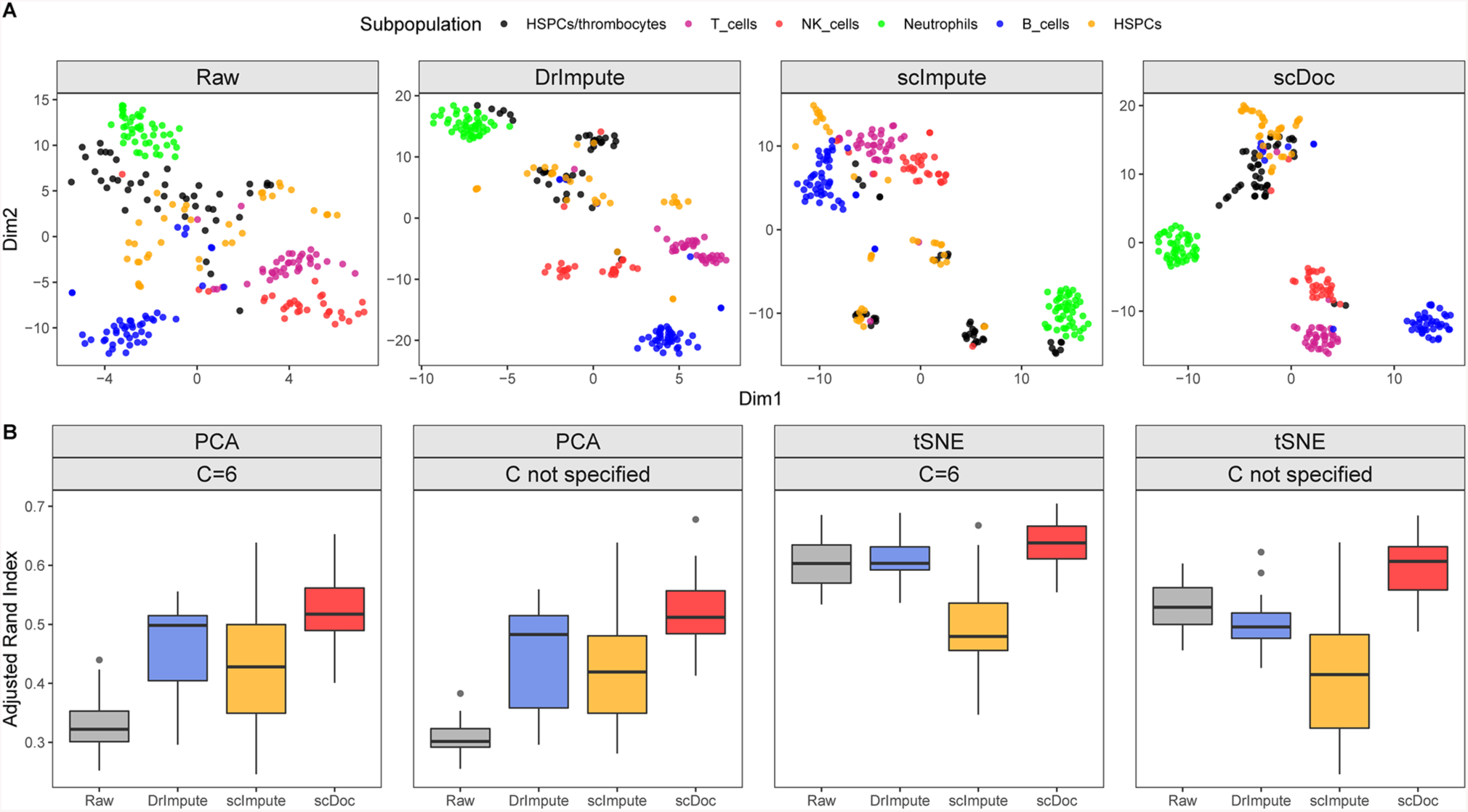
scDoc improved the performance in visualizing zebrafish data and the performance of cell subpopulation identification for the same data. (A) 2D scatterplots using t-SNE. From left to right, panels display the visualization based on the raw and three corrected datasets imputed with DrImpute, scImpute (C=6), and scDoc respectively. (B) The average Adjusted Rand Index of 50 replicates of PCA or t-SNE followed by AP clustering with the true number of subpopulations specified (C=6) or unspecified (C not specified) on the raw and three corrected datasets imputed with DrImpute, scImpute, and scDoc respectively (from left to right).

#### Real dataset 2: lung data

We further demonstrated the capability for scDoc of improving clustering performance in scRNA-seq studies using the lung adenocarcinoma and lung epithelium datasets (Figure 4). Note that the expression profiling of both datasets was given in TPM measurement and the drop-out events were identified using Gamma-Normal mixture model. Same as the previous finding, scDoc improved the performance of visualization using both PCA and t-SNE methods and outperformed other two methods in these two real scRNA-seq studies. In the lung adenocarcinoma study, all three imputation methods improved the performance of visualization for three main different cell types (Figure 4A), “LC.PT.45” (blue and green), “LC.MBT.15” (black), and “H358” (red). But, only t-SNE plot with data imputed by scDoc can group the two batches of “LC.PT.45” together and distinguished these two batches in relatively purer clusters. scDoc was also the only imputation method that can improve the performance of visualization for the two IPSCs cell lines (Figure 4B), BU3 (blue) and ips17 (orange). DrImpute could not help PCA to separate two cell line cells into two groups; while scImpute made more cells go into the opposite cell subpopulation, even though two distinguished groups were observed.

**Figure 4.**
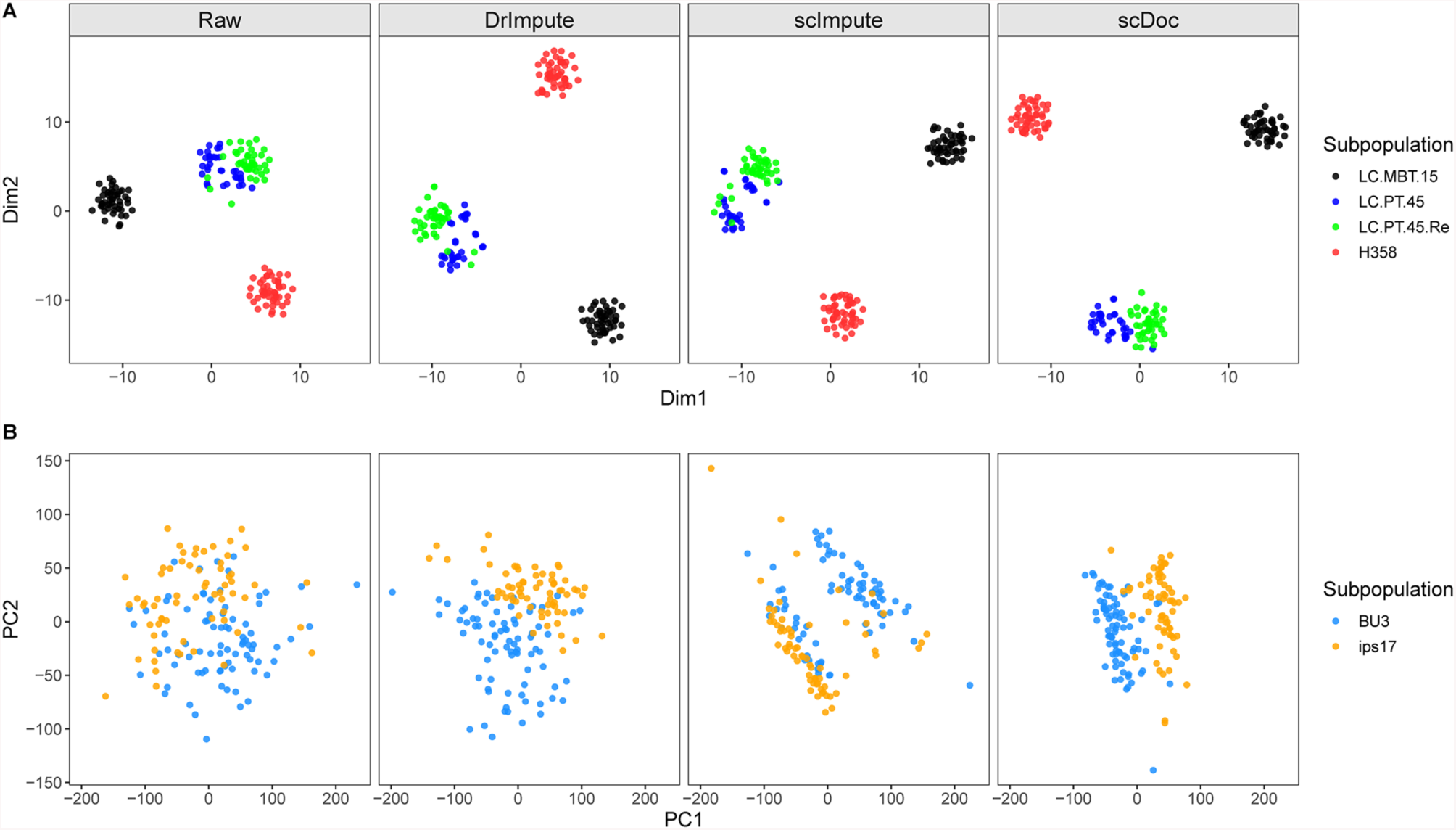
scDoc improved the performance in visualizing lung adenocarcinoma and lung epithelium data. (A) 2D scatterplots of lung adenocarcinoma data using t-SNE. From left to right, panels display the visualization based on the raw and three corrected datasets imputed with DrImpute, scImpute (C=4), and scDoc respectively. (A) 2D scatterplots of lung epithelium data using PCA. From left to right, panels display the visualization based on the raw and three corrected datasets imputed with DrImpute, scImpute (C=2), and scDoc respectively.

In summary, we found that imputing drop-out events in scRNA-seq data with scDoc can significantly improve the performance of visualization for generic dimensionality reduction techniques, such as PCA and t-SNE. In both synthetic and real data analyses, scDoc outperformed two existing imputation methods, DrImpute and scImpute.

### 2.2 scDoc enhances the performance of identification of cell subpopulations

scRNA-seq is widely used to highlight cellular heterogeneity, decompose tissues into cell types, and identify cell subpopulations (Jaitin et al., 2014; Patel et al., 2014; Buettner et al., 2015; Bjorklund et al., 2016). Generic dimensionality reduction techniques (PCA and t-SNE) on selected features followed by clustering methods, e.g. k-means, hierarchical clustering, density-based clustering, and graph clustering, are often applied to scRNA-seq data as a standard protocol (Andrews and Hemberg, 2018). Several R packages including Seurat (Satija et al., 2017), PAGODA (Fan et al., 2016), and SC3 (Kiselev et al., 2017) have been implemented with different combinations of feature selection, dimensionality reduction, and clustering algorithms. In fact, these methods still heavily depend on generic dimensionality reduction techniques that have been proved to be less efficient when drop-outs are present.

To illustrate scDoc’s capacity for improving the performance of identification of cell subpopulations, we used a down-sampling analysis based on the zebrafish dataset. For each of 50 replicates, 80% of 246 cells were randomly picked from the original dataset. We applied dimensionality reduction techniques (PCA and t-SNE) on the raw down-sampled dataset, and down-sampled datasets imputed with scDoc, DrImpute, and scImpute, to extract lower dimensional representatives of datasets. We then ran clustering algorithms on these lower dimensional representatives to partition cells into subpopulations. For the sake of computational intensity, only the first four principal components (PCs) or dimensions (Dims) were taken from PCA or t-SNE and applied to clustering algorithms. We believed the first four PCs or Dims were appropriate choices, because 47% to 63% of the cellular variation can be explained by the first 4 PCs in the down-sampled datasets and cell subpopulations were already distinguishable even in the 2D scatterplot with the first two Dims (Figure 3A). To mimic possible difficulties in real life, we assumed there are two scenarios. In the first one, we know the true number of cell subpopulations as a prior. This assumption is often valid in well-designed experiments where cell types can be identified via marker genes or proteins. In the other scenario, the true number of cell subpopulations is hidden/unknown and shall have to be identified by the clustering algorithm itself. It is common situation for researchers to not have or merely have prior information about the cellular heterogeneity and number of cell subpopulations. To accomplish the comparison under our assumptions, we employed affinity propagation (AP) clustering, which is a clustering algorithm based on the concept of messages passing between data points (Frey and Dueck, 2007). Unlike other clustering algorithms such as k-means, AP clustering does not require us to determine the number of clusters before running the algorithm (Frey and Dueck, 2007). The R version implementation is available in the *apcluster* package (Bodenhofer et al., 2011). We used negative squared Euclidean distance to calculate the similarity matrix, the same as Frey and Dueck demonstrated in the original paper of the AP clustering algorithm. Adjusted Rand Index (ARI) (Hubert and Arabie, 1985) was calculated based on the identified subpopulation labels and their original ones. Higher ARI indicates better agreement between two sets of labels. Figure 3B shows that scDoc dominated performance of identification of cell subpopulations in terms of ARI in all combinations of dimensionality reduction technique and clustering strategies. Further, we note that all three imputation methods improved the performance of PCA followed by AP clustering whenever the true number of subpopulations was given beforehand. However, only scDoc provided significant amelioration of clustering results when applying t-SNE for dimensionality reduction, regardless of whether the number of true subpopulations was specified or not. When using t-SNE plus the AP clustering algorithm, DrImpute showed a minimal improvement when the true number of subpopulations was known, whereas it worsened the performance when it was unspecified. For the data imputed with scImpute, t-SNE followed by AP clustering presented significantly decreased clustering performance compared to that of using the raw down-sampled data. The underperformances of DrImpute and scImpute when using t-SNE as dimensionality reduction, especially when the optimal number of subpopulations was unspecified, were consistent with what we have observed in Figure 2A; DrImpute and scImpute tended to group HSPCs and HSPCs/thrombocytes cells into several small distinct groups. These results suggest that scDoc, accounting drop-outs for cell-to-cell similarity estimation, can be advantageous for generic dimensionality reduction techniques followed by clustering algorithms to identify cell subpopulations in scRNA-seq data.

### 2.3 scDoc boosts the performance of differential expression analysis in scRNA-seq

Differential expression (DE) analysis is one of the most popular tools used in bulk cell RNA-seq data. Though the stochastic expression nature and intra-tissue heterogeneity bring challenges when performing DE analysis in scRNA-seq, they can be compensated at least to some extent because the number of subjects is typically considerably higher in scRNA-seq than in bulk cell RNA-seq (Stegle et al., 2015). Many statistical tools, including the state-of-the-art edgeR and DESeq2, have been developed and widely applied in bulk cell RNA-seq (Anders and Huber, 2010; Robinson et al., 2010). However, these standard RNA-seq DE tools, based on a negative binomial model, suffer in performance due to the presence of drop-out events in scRNA-seq data. Novel methods were recently introduced by using the zero-inflated (SCDE) or hurdle model (MAST).

scDoc was compared with two other scRNA-seq imputation methods (DrImpute and scImpute) for DE analysis, coupling two different standard tools including edgeR and DESeq2, on simulated scRNA-seq datasets. In addition, two scRNA-seq specific DE tools (SCDE and MAST) were applied on the unimputed dataset because these methods include their specific procedures to handle excessive zeros. All these DE tools are implemented and available in Bioconductor packages. Default options were used for all DE tools. Note that the normalization processes were turned off for three standard tools since it was not necessary to adjust the library sizes in synthetic data where true DE effects are known and theoretically would not be affected by the library size. We simulated scRNA-seq datasets for DE analysis (see details in Methods section). Two standard DE tools were applied to both the raw and three imputed datasets. Note that SCDE and MAST were performed on the raw dataset only. Power and family-wise error rate (FWER) of false discovery rate (FDR) adjusted p-values with the Benjamini–Hochberg (BH) procedure (Benjamini and Hochberg, 1995) at a 0.05 significance level were reported to assess the performance of DE analysis. Figure 5A shows that all three scRNA-seq drop-out imputation methods remarkably boosted the performance of all standard DE tools in terms of power compared to the performance in the raw dataset. Further, scDoc significantly outperformed both DrImpute and scImpute for both edgeR and DESeq2. Unsurprisingly, SCDE and MAST, which are designed for zero-inflated datasets, achieved good power for the raw data. However, we also noticed that all three standard DE tools for the data imputed with scDoc resulted in significantly greater power than that of SCDE for the raw dataset. In addition, edgeR and DESeq2 had better DE detection power only for the scDoc imputed dataset than that of MAST for the raw dataset. All standard tools and scRNA-seq specific tools effectively controlled family-wise error rate at a significance level of 0.05. Moreover, we also note that imputation accompanied with standard DE tools was usually less computationally intensive than the scRNA-seq specific DE tools, which need to introduce extra components to model excessive zeros (data not shown). These simulation results suggest that preprocessing scRNA-seq data by imputing drop-outs with scDoc could boost the performance of standard DE tools which are designed for bulk cell RNA-seq. In summary, scDoc outperformed two other imputation methods when working with standard tools. Moreover, scDoc boosted the performance of edgeR and DESeq2 and achieved a better performance than that of using scRNA-seq specific tools in the raw data.

**Figure 5.**
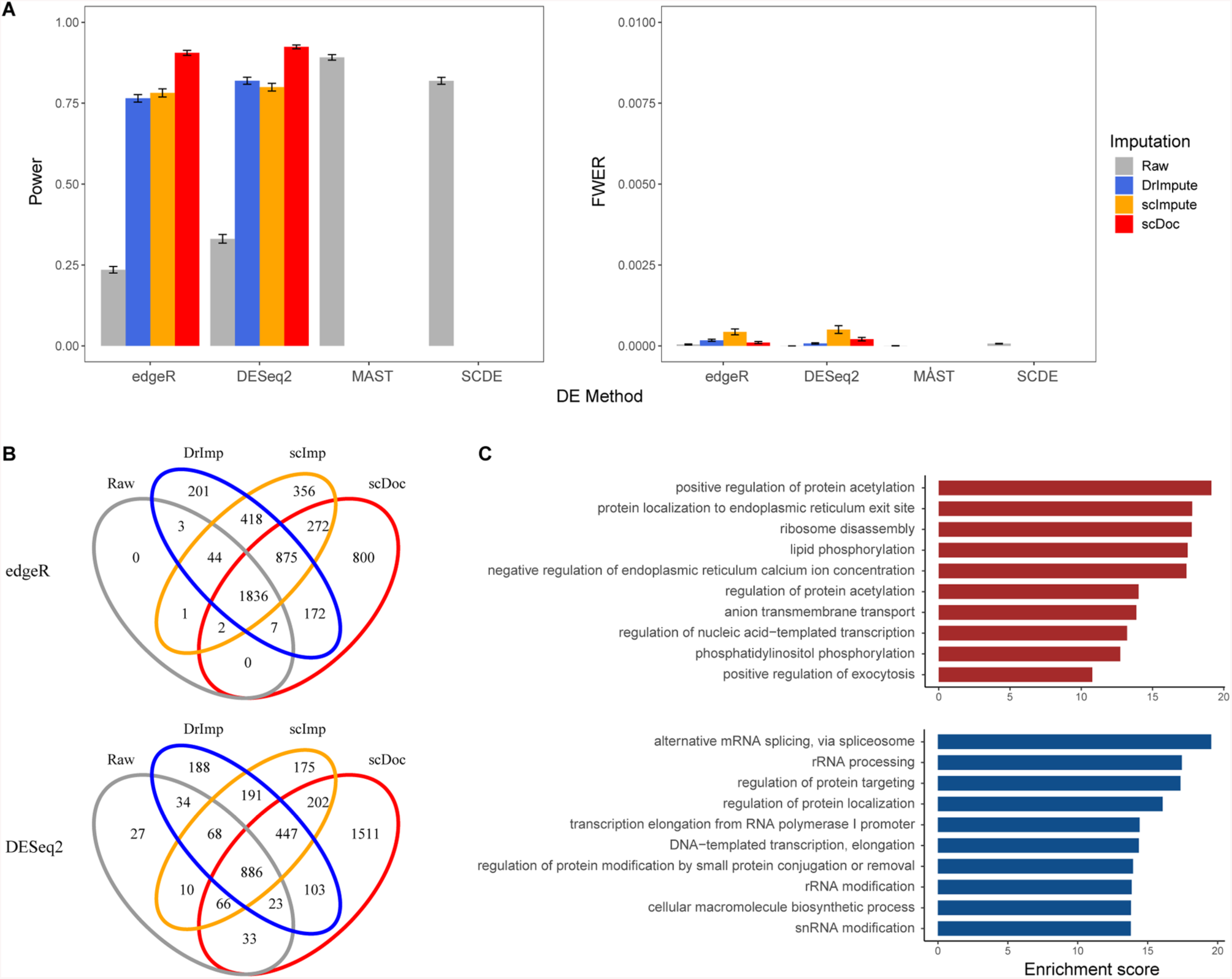
scDoc boosted the performance of the standard DE tools (edgeR and DESeq2). (A) Powers and FWER for detecting DE genes by edgeR and DESeq2 on simulated raw dataset and three corrected datasets imputed with DrImpute, scImpute, and scDoc respectively. Meanwhile, MAST and SCDE are only applied to the raw data. Note that the raw data was introduced excess zeros by using the double exponential function with decay parameter ρ=0.3. (B) Venn diagrams of detected DE gene lists for edgeR (top) and DESeq2 (bottom) applied to the four datasets. (C) Gene ontology (GO) enrichment analysis of unique DE genes by scDoc and edgeR (shown in (B)). Top barplot: top 10 enriched GO terms with upregulated genes in astrocytes; bottom barplot: top 10 enriched GO terms with upregulated genes in oligodendrocytes, from biological process analyzed by *Enrichr* (Kuleshov et al., 2016) are displayed. The enrichment scores calculated by log(p-value x z-score) are used to rank the enrichment of GO terms.

We also demonstrated applications of scDoc to DE detections conducted in the brain data for 62 astrocytes and 32 oligodendrocytes from postnatal cells. After removing lowly expressed genes, 9,317 genes that expressed least 5 counts in least 5 cells were kept in the following DE. Among them, 21 and 23 cell-specific genes were used to validate the DE results for astrocytes and oligodendrocytes, respectively. Before DE analysis, we used visualization methods (PCA and t-SNE) to explore the performance of clustering between astrocytes and oligodendrocytes. Figure S2 (in Appendix) showed that datasets imputed with scDoc presented the most distinct separation between these two cell types. Afterward, two standard DE tools (edgeR and DESeq2) along with their own default normalization methods were applied to the raw and three imputed datasets with scDoc, DrImpute and scImpute respectively. As a result, both edgeR and DESeq2 found the greatest number of DE genes in the dataset imputed with scDoc rather than in the raw or datasets imputed with two other methods (Table 1). Particularly, DESeq2 identified significantly more DE genes in the dataset imputed by scDoc. This is because DESeq2 is designed by default to detect extreme count outliers via Cook’s distance and set the p-value and adjusted p-value of genes with outliers to NA. Due to the advantages of homogeneity within clusters (Figure S2, see Appendix), DESeq2 in the dataset imputed with scDoc detected fewer outliers and gave more robust results. Furthermore, edgeR and DESeq2 in the dataset imputed with scDoc found significantly more DE genes that were not found in the raw or two other imputed datasets (Figure 5B).

**Table 1.** Summary of the number of DE genes detected by edgeR and DESeq2 from the raw the brain dataset (astrocytes and oligodendrocytes only), and three corrected datasets imputed with DrImpute, scImpute (C=2), and scDoc respectively. Note. DE genes were considered significant at FDR < 0.01 and absolute log2FC > 1.5.

|  | Raw | DrImpute | scImpute | scDoc |
| --- | --- | --- | --- | --- |
| **edgeR** | 1893 | 3556 | 3804 | 3964 |
| **DESeq2** | 1147 | 1940 | 2045 | 3271 |

Gene ontology (GO) enrichment analysis showed these additional DE genes are biologically meaningful (Figure 5C). For example, the GO term “positive regulation of exocytosis” is enriched in astrocytes-upregulated genes, which is in agreement with its functions in regulating neurotransmitters and amino acid levels, and releasing neuropeptides and nerontrophins. Additionally, the GO term “regulation of protein targeting” is enriched in oligodendrocytes, which matches its major role as myelination (Ben Haim and Rowitch, 2017; Jana and Pahan, 2017). Moreover, Table 2 showed that edgeR and DESeq2 always found the most cell-specific marker genes for both astrocytes and oligodendrocytes with the data imputed by scDoc. These results from real scRNA-seq data analysis confirmed that scDoc, by imputing drop-out events based on novel cellular similarity assessment metric, could boost the performance of DE analysis for the standard tools originally designed for bulk cell RNA-seq data.

**Table 2.**
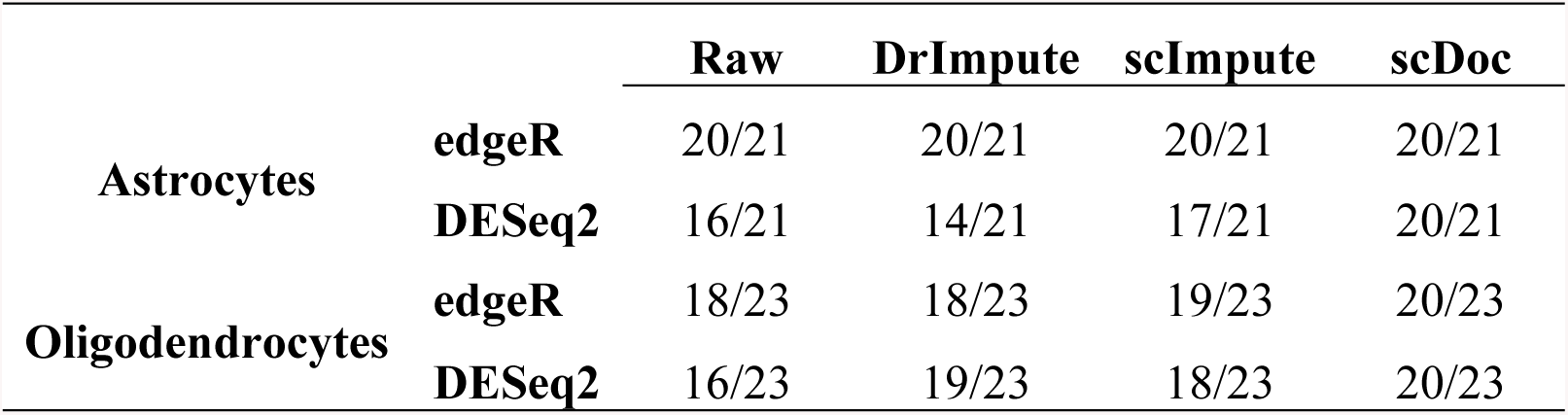
Summary of the number of cell-specific marker genes detected by edgeR and DESeq2 from the raw the brain dataset (astrocytes and oligodendrocytes only), and three corrected datasets imputed with DrImpute, scImpute (C=2), and scDoc respectively. Note: DE genes were considered significant at FDR < 0.01. The marker genes were identified from bulk sequencing of purified cell types in the mouse brain.

## 3 Methods and Data

The flowchart of the proposed method is presented in Figure 1. First, we estimate the drop-out probability for each gene in each individual cell from the input data (raw count) using the PNB mixture model. Second, both input counts and drop-out probability, based on the weighted Cosine similarity metric, are used to estimate cell-to-cell similarity. Third, the detected drop-outs are further evaluated for imputation, which is performed by weighted average from highly similar cells.

### 3.1 Identification of drop-out events

Identification of drop-out events is essential to make the entire imputation accurate and robust. It is necessary to separate technical zeros (due to drop-out) from biological zeros, and technically low expression (near zero counts due to drop-outs) from biologically low expression. Hypothetically, many counts with zeros and near zero values, such as 1 or 2, could be identified as drop-outs; however, a portion of zeros could be considered as biological non-expression values if the overall expression level of that gene across cells is relatively low. Instead of simply treating all zero values as drop-out events or non-zero values as non-drop-outs (like DrImpute), scDoc utilizes a novel two-component mixture model to assess whether an observed count belongs to a drop-out event.

Let *y*_*gi*_ be the read count for gene *g* in cell *i*. For the purpose of notation simplicity, we use *y*_*i*_ to represent the read count for a given gene *g* throughout this subsection. In bulk cell RNA-seq studies, the read count is often assumed to follow a negative binomial (NB) distribution (Anders and Huber, 2010; Robinson et al., 2010),

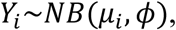

where *μ*_*i*_ is the expected count in sample *i* and *ϕ* is NB dispersion parameter for this gene throughout all available samples. Significantly different from bulk cell RNA-seq, scRNA-seq has many zeros and near zero counts, known as “drop-outs”, which cannot be handled merely by the previous NB model. Thus, spectacular tools are required to effectively model gene expression while accounting for drop-outs. Due to its flexibility, the mixture model becomes a potential solution for zero-inflated data sets. Recently, several methods have been proposed based on mixture models such as the Zero-inflated Negative Binomial model (Ran and Daye, 2017; Van den Berge et al., 2018), the Gamma-Normal mixture model (Li and Li, 2018), and a three-component mixture model (Kharchenko et al., 2014). Here, we propose a Poisson-Negative binomial (PNB) model in which the first component, Poisson distribution, is used to account for the “drop-outs”, and the second component, negative binomial distribution, is used to represent the “actual expression” (it could be zero or a non-zero value). The PNB formulation has the density function,

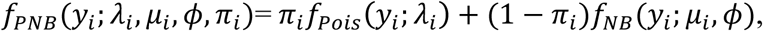

where *π*_*i*_ is the probability that an observed count belongs to a drop-out event for that gene in cell *i, λ*_*i*_ is the mean parameter of Poisson distribution, and *μ*_*i*_ and *ϕ* are the mean and dispersion parameters of NB distribution as described previously.

All these parameters can be estimated via maximum likelihood (ML) estimations with an expectation maximization (EM) algorithm (Dempster et al., 1977). The posterior probability *d*_*gi*_ (or *d*_*i*_ for simplicity) is then calculated as:

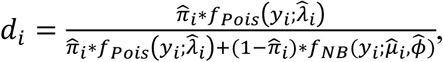

where 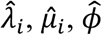, and 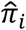 were the estimates from the above EM algorithm. This probability measures how likely an observed count of gene *g* in cell *i* belongs to drop-outs. Throughout both the ‘Methods and Data’ and ‘Results’ sections, any observed count with drop-out probability greater than 0.5 will be considered a drop-out event to be imputed. The drop-out rate for a gene is defined as the proportion of detected drop-out events across all cells.

Begin with raw counts brings lots of flexibility and accuracy, however, TPM (Transcripts Per kilobase Million) which normalizes not only gene length but also sequencing depth, is now becoming popular and the default output formatting in several popular RNA-seq alignment software. TPM, as continuous measurements, cannot work with the introduced Poisson-NB mixture model and requires other solution in this drop-out identification step. Thus, we adopted the Gamma-Normal mixture model, which was introduced in the scImpute (Li and Li, 2018), as an alternative to deal with logarithm transformed TPM measurements. Note that normalizing expression matrix by the library size of each sample is skipped, because TPM measurements have been normalized. Like the Poisson-NB mixture model, the Gamma-Normal mixture uses the two components (Gamma and Normal) to take account for “drop-out” and “actual expression”, respectively. However, we noticed that the “Gamma” component could not always fit the drop-out part well in real scRNA-seq data, especially the proportion of zeros is high. Here, as a solution on such extreme occasion, we modified the method and substituted Exponential distribution for the “Gamma” component. After all parameters were estimated from EM algorithm, the posterior can be obtained to represent the drop-out probability of gene *g* in cell *i*. Using the predefined threshold, any observed expression value with greater drop-out probability will be considered as a drop-out event to be imputed in the following steps.

### 3.2 Calculation of cell-to-cell similarity

scDoc is designed to impute the identified drop-out events by borrowing information from highly similar cells. Taking advantage of including drop-outs in the cell-to-cell similarity assessment, we introduce a new similarity metric, Weighted Cosine similarity (WCS), which is based on both the drop-out probability and Cosine similarity. The Cosine similarity is a measure of similarity between two non-zero vectors of an inner product space that measures the cosine of the angle between them and is widely used in text mining for sparse vectors (Singhal, 2001). Given two *m*-dimensional non-zero vectors, A and B, the Cosine similarity, cos(θ), is defined as

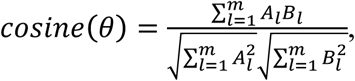

where *A*_*l*_ and *B*_*l*_ are features in vector *A* and *B*, respectively.

The weighted cosine similarity between cell *i* and *j* across all *p* genes is defined as

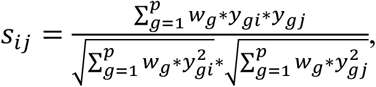

where *w*_*g*_ = 1 when both of *y*_*gi*_ and *y*_*gj*_ are non-drop-out values or drop-out events; otherwise, we call this pair of data a “questionable” pair and let *w*_*g*_ equal the drop-out rate for this gene as the gene-specific weight. In other words, WCS assigns weights with respect to how “confident” we are with each pair of observed counts (between two cells) contributing to the cell similarity. For instance, a “questionable” pair from a gene with low drop-out rate will be assigned a small weight (which results in a greater *s*), whereas a large weight (i.e., a smaller *s*) will be assigned when drop-out rate is high. The intuition behind the choice is that drop-out events are more likely to have greater influences on those genes that have high expressions and few drop-out events when assessing cell-to-cell similarities. To understand how WCS works, assume two observed counts from cell *i* and *j* for gene *g* are measured. Hypothetically, there could be three different situations in terms of their drop-out probabilities; (1) neither of the counts are drop-out events, i.e., *d*_*i*_ ≤ 0.5 and *d*_*j*_ ≤ 0.5, (2) both counts are drop-out events, *d*_*i*_ > 0.5 and *d*_*j*_ > 0.5, (3) only one of the counts is drop-out event, either *d*_*i*_ > 0.5 or *d*_*j*_ > 0.5. For the first two situations, there is no need to put a smaller weight on that pair when calculating similarity, because either we are confident with both measurements or that pair of drop-out values has minimal contributions to the calculation. In the last scenario, however, that pair of observations would incorrectly exaggerate the difference between two cells when they in fact belong to the same cell subpopulation, thus a bias would be introduced, and cell-to-cell similarities be distorted. Therefore, we give less weight for the “questionable” pair of data in the similarity calculation. Theoretically, *w* can be any value between 0 and 1, where 0 and 1 are two extreme cases; 0 means that we totally remove that “questionable” pair of data from calculation, 1 means that we completely ignore the drop-out nature and use the original Cosine similarity. Any value between 0 and 1 indicates that we keep that pair of data but understate them in similarity estimation. After calculating the similarity between any pair of cells, we can obtain a similarity matrix that is used to represent the cell-to-cell similarity. Note that, before we compute the WCS, the expression count matrix is logarithm transformed along with adding one pseudo count.

Different from all existing imputation methods in scRNA-seq data, scDoc will be the first imputation method taking the detected drop-outs into account when evaluating cell-to-cell similarities. In doing this, it can identify highly similar cells accurately as a result of eliminating the unignorable influence of drop-outs, which has been indicated to distort the true cell-to-cell relationship (Lin et al., 2017).

### 3.3 Imputation of drop-out values

For each identified drop-out event, scDoc imputes the drop-out values by borrowing information, the expression values in the same gene, from other highly similar cells. The general imputation strategy for each gene *g* is described as follows:

1. Find *k* highly similar cells to a target cell (i.e., in which gene *g* has (a) drop-out events detected) according to the calculated cell-to-cell similarity between all other cells and the target one. The default number for *k* is 6.
2. Compare the similarity between the most similar one among these *k* candidate cells and the target cell to a pre-defined threshold.
  - If the most similar candidate fails to pass the threshold, this cell will be detected and reported as a potential outlier and drop-out values will not be imputed (this detection is off as default in scDoc package).
  - Otherwise, the drop-out value will be considered further in next step.
3. If the target cell is not detected as an outlier, scDoc makes another judgement based on cell-to-cell similarities to decide whether it would impute the drop-out value for gene *g* in cell *i*. The final imputation procedure can be described as:
  - If there are two or more candidate cells having non-drop-out values, the drop-out value will be replaced by the weighted average,

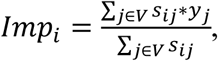
  - where *s*_*ij*_ is the weighted Cosine similarity between the target cell *i* and candidate cell *j*, and *V* is a set of indexes for all candidate cells.
  - If there is only one candidate cell having a non-drop-out value for that gene, the drop-out value will be imputed based upon the WCS measure. For instance, if the candidate cell has a non-drop-out value and its WCS similarity is greater than 90^th^ percentile of the overall cell-to-cell similarities, the drop-out value will be replaced by the expression value in that candidate cell. If the 90^th^ percentile criterion is not met, the drop-out event will not be imputed.
  - If all expression values of the candidate cells are drop-out events, no imputation is needed for that gene in the target cell.

Steps 1-3 will be applied to each gene. Note that, based on the above strategy, scDoc will only replace some zeros with a nonzero value when the drop-out probability is above 0.5; and it will correct some low expression as well. In other words, this imputation strategy not only protects the scRNA-seq data from being over-imputed for zero counts but also could correct some low count values.

### 3.4 Real scRNA-seq Datasets and data preprocessing

To evaluate the performance of the new method, we consider the datasets from real scRNA-seq studies. One dataset (referred to as ‘Zebrafish’ dataset) (Tang et al., 2017) can be obtained from Gene Expression Omnibus (GEO) with accession number GSE100911. It established the transcriptional profiles of 246 single cells of whole kidney marrow and thymic T cells isolated from adult, transgenic zebrafish using FACS. The cells were sorted and labeled into six groups including HSPCs (n=39), HSPCs/thrombocytes (n=43), neutrophils (n=47), T cells (n=38), NK cells (n=31), and B cells (n=48). Note that HSPCs and HSPCs/thrombocytes, though labeled by different markers, were from one defined cell group with expected heterogeneity. In this study we treated them as two distinct groups, though they are difficult to separate clearly.

The human brain dataset (Darmanis et al., 2015) can be obtained from GEO with accession number GSE67835. This scRNA-seq study sequenced 466 cells from human cortical tissue including cell types: oligodendrocytes (n=38), astrocytes (n=62), microglia (n=16), neurons (n=131), endothelial (n=20), fetal quiescent (n=110), and fetal replicating cells (n=25). This dataset also provided a list of cell-specific marker genes that are identified from bulk sequencing of purified cell types in the mouse brain (Zhang et al., 2014).

The expression matrix of TPM measurements for the lung adenocarcinoma dataset and the lung epithelium dataset were obtained from GEO accessions GSE69405 and GSE96106, respectively. The lung adenocarcinoma dataset (Kim et al., 2015) contains transcriptome profiles of single cancer cells originating from lung adenocarcinoma patient-derived xenograft (PDX) tumors. It includes PDX cells (LC-PT-45, n=34), PDX cells additional batch (LC-Pt-45-Re, n=43), H358 human lung cancer cells (H358, n=50) as cell line control, and another lung cancer PDX case (LC-MBT-15, n=49) as validation. The lung epithelium dataset (Hawkins et al., 2017) provides the scRNA-seq profiling of single lung epithelial cells from human induced pluripotent stem cells (IPSCs). Single cells were collected from two cell lines BU3 (n=80) and ips17 (65) on day 15 of the lung directed differentiation and loaded into separate Fluidigm C1 integrated fluidics circuits (IFCs).

Filtering out lowly expressed genes and low-quality cells is optional; however, it could help reduce noise and improve imputation accuracy. We applied a general gene filtering process on all the datasets used throughout analyses. Particularly, only genes expressing at least 5 counts in 5 or more cells were kept for further steps. We also notice that the filtering threshold on the number of cells should be chosen to accommodate the potential group size (i.e., number of cells in the group) in the smallest subpopulation. Low quality cells with extremely low total counts (library size) are also suggested to be removed according to the distribution of total counts in each study. However, we did not apply the cell quality filtering for the two datasets used in the analyses since they have been well checked and documented.

## 4 Discussion

In this paper, we have demonstrated that scDoc offers a robust and accurate imputation method for correcting drop-out events in scRNA-seq data. scDoc utilizes a novel Poisson-Negative Binomial mixture where the drop-outs and actual expression values are modeled via two different components separately. The model also takes the cell library sizes into account, and then computes the drop-out probability for a gene in each cell. Trying to avoid over-imputing the expression measurements (which might not be affected by drop-out events) and changing the stochastic nature of gene expression, scDoc only imputes the identified drop-out events, the observed zero or near zero counts having high probability for being dropped. scDoc is the first imputation method that accounts for drop-outs when assessing cell-to-cell similarity, which is the crucial step before imputation. As drop-out events have non-neglectful influences on the generic dimensionality reduction techniques, cellular similarity estimates based on these techniques are distorted. Drop-out events imputation that uses the biased cell-to-cell relationship would misrepresent the true signal of biological interests. To overcome the challenge of investigating the inter-cell similarity present in drop-outs, scDoc develops a novel approach in which the cell-to-cell similarity is estimated by weighted Cosine similarity, which modifies the traditional Cosine similarity in order to incorporate gene-specific weight according to the drop-out rate of that gene. Finally, the robust imputation procedure based on both drop-out probability and cellular similarity is proposed for replacing drop-out values. Through comprehensive simulation studies, scDoc has been shown advantageous for the analysis of data visualization, cell subpopulation clustering, and differential expression analysis in scRNA-seq data. Furthermore, scDoc can improve the performance of generic dimensionality reduction techniques and standard DE tools. Applications of scDoc to the analyses of the zebrafish dataset illustrate that preprocessing the scRNA-seq dataset by imputing drop-out events with scDoc would help not only identify the most clearly distinct subpopulations but also correctly discover the heterogeneity within a cell subpopulation.

Our proposed weighted Cosine similarity in scDoc is related to the soft Cosine similarity, which considers the features in the non-zero vectors as dependent, while the traditional Cosine similarity treats them as independent. In soft Cosine similarity, the concept of cosine is generalized by considering the similarity/correlation of features, which are used in the calculation as weight (Sidorov et al., 2012). In this paper, we used the weight-based drop-out probabilities of a pair of observations. We noted that this weight can be analogously derived from the relationship between tw oexpression values in terms of the property of drop-outs. In the three types of relationships for an observed pair of counts in a cell, the similarity of the two observations in this pair is 1 in terms of drop-out probability when both or neither of them are identified as drop-out events, while the similarity is equal to the drop-out rate of this cell when only one of them is a drop-out event. Note that this soft Cosine similarity, explained by relationship in terms of drop-out, is equivalent to our proposed weighted Cosine similarity, though derived from distinct thought processes in relation to the drop-out nature in scRNA-seq data.

scDoc identifies potential drop-out events with a PNB mixture model, rather than merely considering all zero expressions as drop-outs and all non-zero measurements as actual expression values. It brings the advantages of an efficient elimination of influence from drop-out events and a prevention of over-imputing on the true non-expression values. Estimates in the PNB model are obtained by employing the E-M algorithm, though this brings a trade-off between accuracy and complexity. To make sure scDoc is acceptable in terms of computational intensity, we provide gradients to the ‘BFGS’ optimization algorithm and make parallel processing available for both Windows and Unix users.

Drop-out detection in the visualization simulation studies for 15,000 genes in 150 cells took about 3 minutes using a single process on a Windows laptop with 2.60 GHz Intel i5-7300U CPUs and 16GB RAM, and R (version 3.4.3, 64-bit). In real data analysis using the same computation platform, it took ∼8 mins for 11,307 genes and 246 cells in the zebrafish dataset and ∼18 mins for 9,317 genes and 94 cells in the brain dataset. Though we believe it is worth the accuracy for the price of a slight increase in computational intensity, the efficacy may be improved by using a more efficient optimization method to estimate drop-out probabilities.

### Appendix

**Figure S1.**
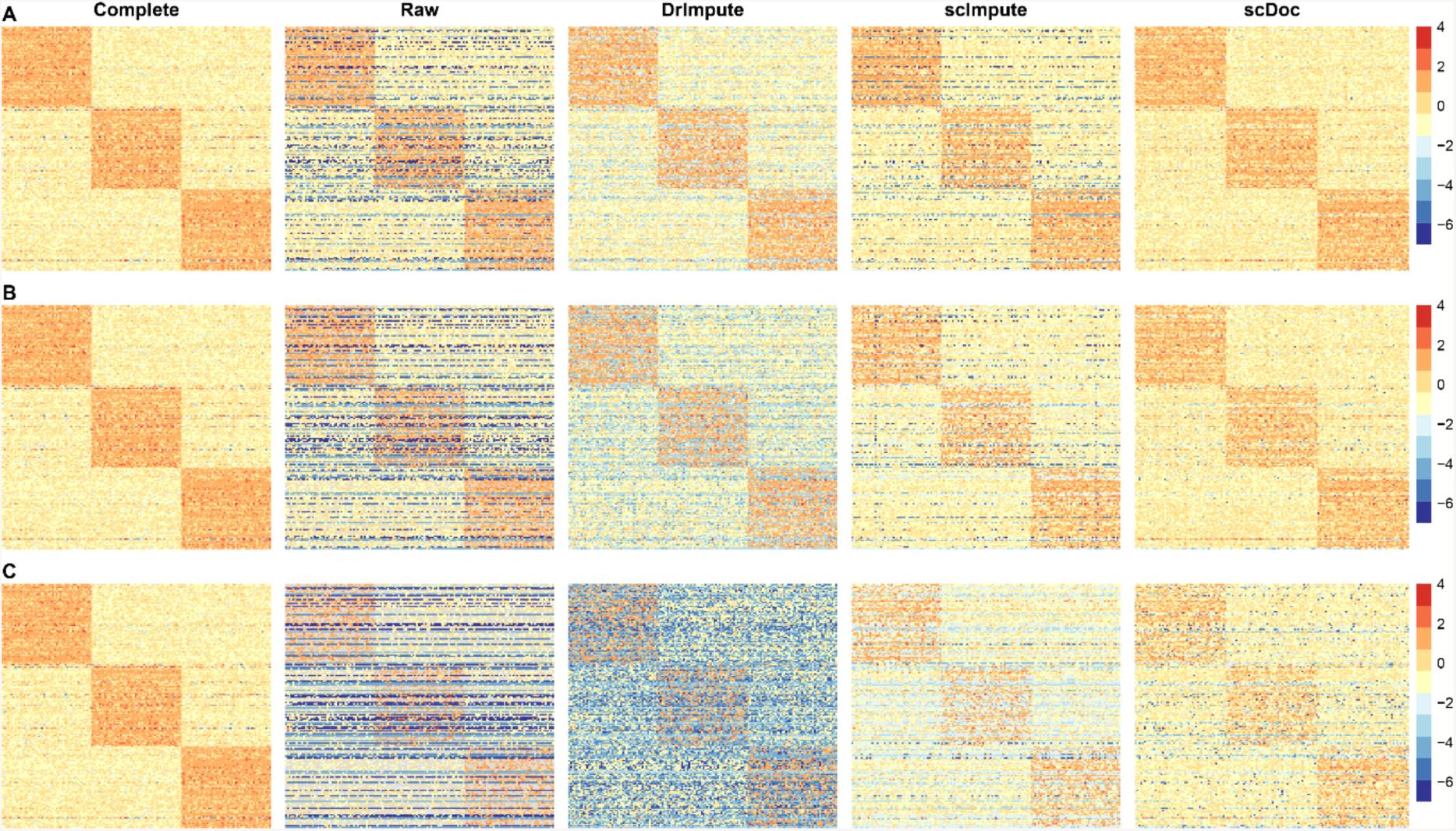
scDoc corrected drop-out events and improved the visualization of simulated scRNA-seq data. Heatmaps of the expression profiles of 180 DE genes in the complete and raw, and three corrected datasets imputed with DrImpute, scImpute (C=3), and scDoc respectively (from left to right). Excessive zeros were introduced to the raw dataset using the double exponential function with decay parameter *ρ*. (A) *ρ* = 0.3. The zero prevalence is ∼32%. (B) *ρ* = 0.2. The zero prevalence is ∼43%. (C) *ρ* = 0.1. The zero prevalence is ∼62%.

**Figure S2.**
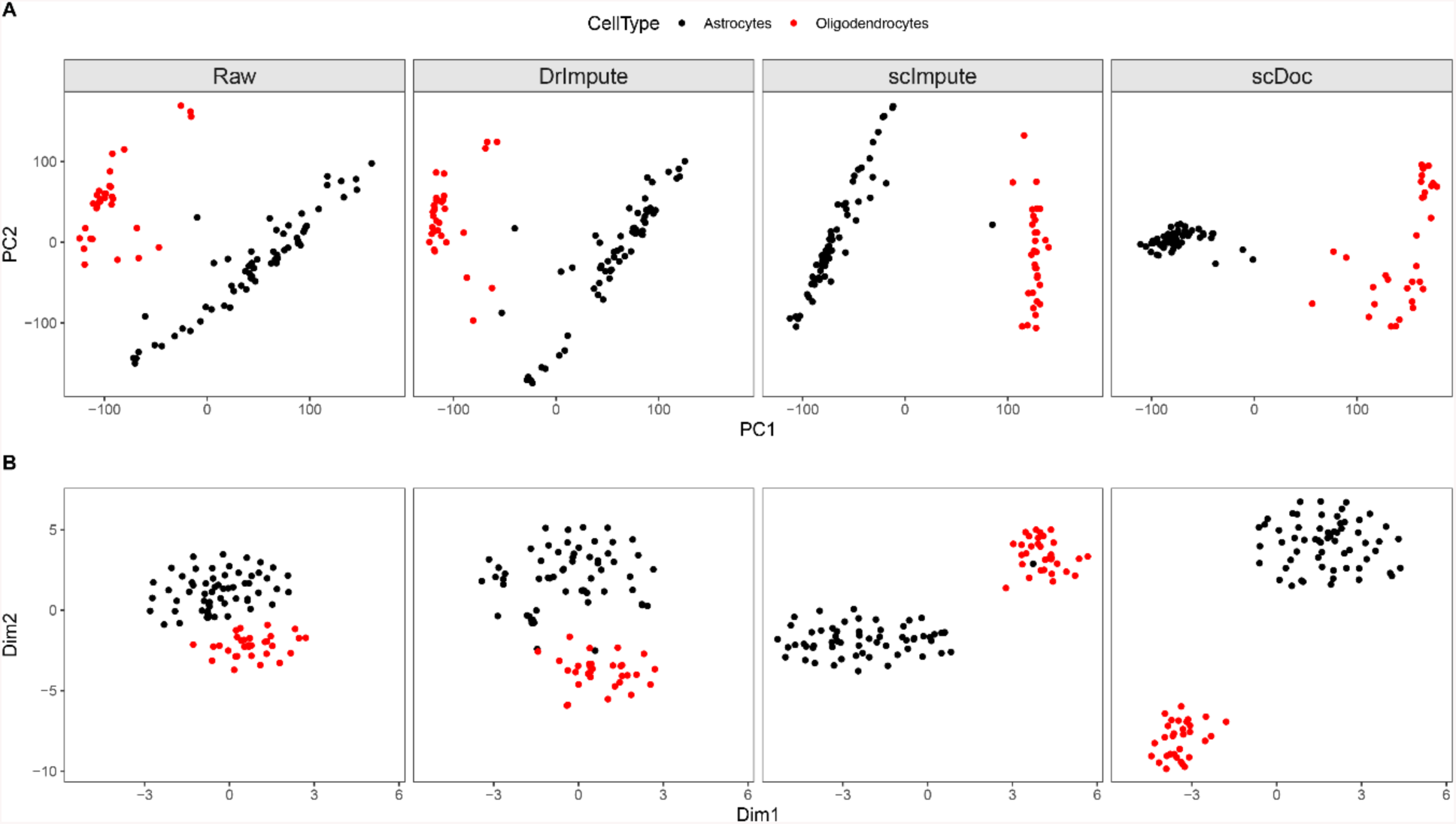
scDoc improved the performance of visualizing the brain scRNA-seq data. (A) 2D plots of the first two PCs calculated from the raw dataset (astrocytes and oligodendrocytes only), and three corrected datasets imputed with DrImpute, scImpute (C=2), and scDoc respectively (from left to right). (B) 2D visualizations using t-SNE on the same four datasets in (A) (from left to right).

## References

Anders, S., and Huber, W. (2010). Differential expression analysis for sequence count data. Genome Biol 11(10), R106. doi: 10.1186/gb-2010-11-10-r106.

Andrews, T.S., and Hemberg, M. (2018). Identifying cell populations with scRNASeq. Mol Aspects Med 59, 114–122. doi: 10.1016/j.mam.2017.07.002.

Bacher, R., and Kendziorski, C. (2016). Design and computational analysis of single-cell RNA-sequencing experiments. Genome Biol 17, 63. doi: 10.1186/s13059-016-0927-y.

Ben Haim, L., and Rowitch, D.H. (2017). Functional diversity of astrocytes in neural circuit regulation. Nat Rev Neurosci 18(1), 31–41. doi: 10.1038/nrn.2016.159.

Benjamini, Y., and Hochberg, Y. (1995). Controlling the False Discovery Rate - a Practical and Powerful Approach to Multiple Testing. Journal of the Royal Statistical Society Series B-Methodological 57(1), 289–300.

Bjorklund, A.K., Forkel, M., Picelli, S., Konya, V., Theorell, J., Friberg, D., et al. (2016). The heterogeneity of human CD127(+) innate lymphoid cells revealed by single-cell RNA sequencing. Nat Immunol 17(4), 451–460. doi: 10.1038/ni.3368.

Blakeley, P., Fogarty, N.M., Del Valle, I., Wamaitha, S.E., Hu, T.X., Elder, K., et al. (2015). Defining the three cell lineages of the human blastocyst by single-cell RNA-seq. Development 142(20), 3613. doi: 10.1242/dev.131235.

Bodenhofer, U., Kothmeier, A., and Hochreiter, S. (2011). APCluster: an R package for affinity propagation clustering. Bioinformatics 27(17), 2463–2464. doi: 10.1093/bioinformatics/btr406.

Buettner, F., Natarajan, K.N., Casale, F.P., Proserpio, V., Scialdone, A., Theis, F.J., et al. (2015). Computational analysis of cell-to-cell heterogeneity in single-cell RNA-sequencing data reveals hidden subpopulations of cells. Nat Biotechnol 33(2), 155–160. doi: 10.1038/nbt.3102.

Darmanis, S., Sloan, S.A., Zhang, Y., Enge, M., Caneda, C., Shuer, L.M., et al. (2015). A survey of human brain transcriptome diversity at the single cell level. Proc Natl Acad Sci U S A 112(23), 7285–7290. doi: 10.1073/pnas.1507125112.

Dempster, A.P., Laird, N.M., and Rubin, D.B. (1977). Maximum Likelihood from Incomplete Data via the EM Algorithm. Journal of the Royal Statistical Society. Series B (Methodological) 39(1), 1–38.

Deng, Q., Ramskold, D., Reinius, B., and Sandberg, R. (2014). Single-cell RNA-seq reveals dynamic, random monoallelic gene expression in mammalian cells. Science 343(6167), 193–196. doi: 10.1126/science.1245316.

Fan, J., Salathia, N., Liu, R., Kaeser, G.E., Yung, Y.C., Herman, J.L., et al. (2016). Characterizing transcriptional heterogeneity through pathway and gene set overdispersion analysis. Nature Methods 13(3), 241-+. doi: 10.1038/Nmeth.3734.

Finak, G., McDavid, A., Yajima, M., Deng, J., Gersuk, V., Shalek, A.K., et al. (2015). MAST: a flexible statistical framework for assessing transcriptional changes and characterizing heterogeneity in single-cell RNA sequencing data. Genome Biol 16, 278. doi: 10.1186/s13059-015-0844-5.

Frey, B.J., and Dueck, D. (2007). Clustering by passing messages between data points. Science 315(5814), 972–976. doi: 10.1126/science.1136800.

Gong, W., Kwak, I.Y., Pota, P., Koyano-Nakagawa, N., and Garry, D.J. (2018). DrImpute: imputing dropout events in single cell RNA sequencing data. BMC Bioinformatics 19(1), 220. doi: 10.1186/s12859-018-2226-y.

Han, Y., Gao, S., Muegge, K., Zhang, W., and Zhou, B. (2015). Advanced Applications of RNA Sequencing and Challenges. Bioinform Biol Insights 9(Suppl 1), 29–46. doi: 10.4137/BBI.S28991.

Hawkins, F., Kramer, P., Jacob, A., Driver, I., Thomas, D.C., McCauley, K.B., et al. (2017). Prospective isolation of NKX2-1-expressing human lung progenitors derived from pluripotent stem cells. J Clin Invest 127(6), 2277–2294. doi: 10.1172/JCI89950.

Huang, M., Wang, J., Torre, E., Dueck, H., Shaffer, S., Bonasio, R., et al. (2018). SAVER: gene expression recovery for single-cell RNA sequencing. Nat Methods 15(7), 539–542. doi: 10.1038/s41592-018-0033-z.

Hubert, L., and Arabie, P. (1985). Comparing Partitions. Journal of Classification 2(2-3), 193–218. doi: Doi 10.1007/Bf01908075.

Jaitin, D.A., Kenigsberg, E., Keren-Shaul, H., Elefant, N., Paul, F., Zaretsky, I., et al. (2014). Massively parallel single-cell RNA-seq for marker-free decomposition of tissues into cell types. Science 343(6172), 776–779. doi: 10.1126/science.1247651.

Jana, M., and Pahan, K. (2017). Astrocytes, Oligodendrocytes and Schwann Cells. 117–140. doi: 10.1007/978-3-319-44022-4_10.

Kharchenko, P.V., Silberstein, L., and Scadden, D.T. (2014). Bayesian approach to single-cell differential expression analysis. Nat Methods 11(7), 740–742. doi: 10.1038/nmeth.2967.

Kim, K.T., Lee, H.W., Lee, H.O., Kim, S.C., Seo, Y.J., Chung, W., et al. (2015). Single-cell mRNA sequencing identifies subclonal heterogeneity in anti-cancer drug responses of lung adenocarcinoma cells. Genome Biol 16, 127. doi: 10.1186/s13059-015-0692-3.

Kiselev, V.Y., Kirschner, K., Schaub, M.T., Andrews, T., Yiu, A., Chandra, T., et al. (2017). SC3: consensus clustering of single-cell RNA-seq data. Nat Methods 14(5), 483–486. doi: 10.1038/nmeth.4236.

Kuleshov, M.V., Jones, M.R., Rouillard, A.D., Fernandez, N.F., Duan, Q., Wang, Z., et al. (2016). Enrichr: a comprehensive gene set enrichment analysis web server 2016 update. Nucleic Acids Res 44(W1), W90–97. doi: 10.1093/nar/gkw377.

Li, W.V., and Li, J.J. (2018). An accurate and robust imputation method scImpute for single-cell RNA-seq data. Nat Commun 9(1), 997. doi: 10.1038/s41467-018-03405-7.

Lin, P., Troup, M., and Ho, J.W. (2017). CIDR: Ultrafast and accurate clustering through imputation for single-cell RNA-seq data. Genome Biol 18(1), 59. doi: 10.1186/s13059-017-1188-0.

Lun, A.T., Bach, K., and Marioni, J.C. (2016). Pooling across cells to normalize single-cell RNA sequencing data with many zero counts. Genome Biol 17, 75. doi: 10.1186/s13059-016-0947-7.

Moussa, M., and Mandoiu, I. (2019). Locality Sensitive Imputation for Single-Cell RNA-Seq Data. Journal of Computational Biology. https://doi.org/10.1089/cmb.2018.0236.

Patel, A.P., Tirosh, I., Trombetta, J.J., Shalek, A.K., Gillespie, S.M., Wakimoto, H., et al. (2014). Single-cell RNA-seq highlights intratumoral heterogeneity in primary glioblastoma. Science 344(6190), 1396–1401. doi: 10.1126/science.1254257.

Petropoulos, S., Edsgard, D., Reinius, B., Deng, Q., Panula, S.P., Codeluppi, S., et al. (2016). Single-Cell RNA-Seq Reveals Lineage and X Chromosome Dynamics in Human Preimplantation Embryos. Cell 167(1), 285. doi: 10.1016/j.cell.2016.08.009.

Pierson, E., and Yau, C. (2015). ZIFA: Dimensionality reduction for zero-inflated single-cell gene expression analysis. Genome Biol 16, 241. doi: 10.1186/s13059-015-0805-z.

Poulin, J.F., Tasic, B., Hjerling-Leffler, J., Trimarchi, J.M., and Awatramani, R. (2016). Disentangling neural cell diversity using single-cell transcriptomics. Nat Neurosci 19(9), 1131–1141. doi: 10.1038/nn.4366.

Ran, D., and Daye, Z.J. (2017). Gene expression variability and the analysis of large-scale RNA-seq studies with the MDSeq. Nucleic Acids Res 45(13), e127. doi: 10.1093/nar/gkx456.

Risso, D., Perraudeau, F., Gribkova, S., Dudoit, S., and Vert, J.P. (2018). A general and flexible method for signal extraction from single-cell RNA-seq data. Nat Commun 9(1), 284. doi: 10.1038/s41467-017-02554-5.

Robinson, M.D., McCarthy, D.J., and Smyth, G.K. (2010). edgeR: a Bioconductor package for differential expression analysis of digital gene expression data. Bioinformatics 26(1), 139–140. doi: 10.1093/bioinformatics/btp616.

Satija, R., Butler, A., and Hoffman, P. (2017). “Seurat: Tools for Single Cell Genomics”. R package version).

Shalek, A.K., Satija, R., Adiconis, X., Gertner, R.S., Gaublomme, J.T., Raychowdhury, R., et al. (2013). Single-cell transcriptomics reveals bimodality in expression and splicing in immune cells. Nature 498(7453), 236–240. doi: 10.1038/nature12172.

Sidorov, G., Velasquez, F., Stamatatos, E., Gelbukh, A., and Chanona-Hernández, L. (Year). “Syntactic dependency-based n-grams as classification features”, in: Mexican International Conference on Artificial Intelligence: Springer), 1–11.

Singhal, A. (2001). Modern Information Retrieval: A Brief Overview. Bulletin of the IEEE Computer Society Technical Committee on Data Engineering 24(4), 35–43.

Stegle, O., Teichmann, S.A., and Marioni, J.C. (2015). Computational and analytical challenges in single-cell transcriptomics. Nat Rev Genet 16(3), 133–145. doi: 10.1038/nrg3833.

Tang, F., Barbacioru, C., Bao, S., Lee, C., Nordman, E., Wang, X., et al. (2010). Tracing the derivation of embryonic stem cells from the inner cell mass by single-cell RNA-Seq analysis. Cell Stem Cell 6(5), 468–478. doi: 10.1016/j.stem.2010.03.015.

Tang, Q., Iyer, S., Lobbardi, R., Moore, J.C., Chen, H., Lareau, C., et al. (2017). Dissecting hematopoietic and renal cell heterogeneity in adult zebrafish at single-cell resolution using RNA sequencing. J Exp Med 214(10), 2875–2887. doi: 10.1084/jem.20170976.

Trapnell, C., Cacchiarelli, D., Grimsby, J., Pokharel, P., Li, S., Morse, M., et al. (2014). The dynamics and regulators of cell fate decisions are revealed by pseudotemporal ordering of single cells. Nat Biotechnol 32(4), 381–386. doi: 10.1038/nbt.2859.

Treutlein, B., Brownfield, D.G., Wu, A.R., Neff, N.F., Mantalas, G.L., Espinoza, F.H., et al. (2014). Reconstructing lineage hierarchies of the distal lung epithelium using single-cell RNA-seq. Nature 509(7500), 371–375. doi: 10.1038/nature13173.

Van den Berge, K., Perraudeau, F., Soneson, C., Love, M.I., Risso, D., Vert, J.P., et al. (2018). Observation weights unlock bulk RNA-seq tools for zero inflation and single-cell applications. Genome Biol 19(1), 24. doi: 10.1186/s13059-018-1406-4.

van Dijk, D., Nainys, J., Sharma, R., Kathail, P., Carr, A.J., Moon, K.R., et al. (2017). MAGIC: A diffusion-based imputation method reveals gene-gene interactions in single-cell RNA-sequencing data. bioRxiv. doi: 10.1101/111591.

Villani, A.C., Satija, R., Reynolds, G., Sarkizova, S., Shekhar, K., Fletcher, J., et al. (2017). Single-cell RNA-seq reveals new types of human blood dendritic cells, monocytes, and progenitors. Science 356(6335). doi: 10.1126/science.aah4573.

Wang, Z., Gerstein, M., and Snyder, M. (2009). RNA-Seq: a revolutionary tool for transcriptomics. Nat Rev Genet 10(1), 57–63. doi: 10.1038/nrg2484.

Wu, Z., Zhang, Y., Stitzel, M.L., and Wu, H. (2018). Two-phase differential expression analysis for single cell RNA-seq. Bioinformatics. doi: 10.1093/bioinformatics/bty329.

Xue, Z., Huang, K., Cai, C., Cai, L., Jiang, C.Y., Feng, Y., et al. (2013). Genetic programs in human and mouse early embryos revealed by single-cell RNA sequencing. Nature 500(7464), 593–597. doi: 10.1038/nature12364.

Zhang, Y., Chen, K., Sloan, S.A., Bennett, M.L., Scholze, A.R., O’Keeffe, S., et al. (2014). An RNA-sequencing transcriptome and splicing database of glia, neurons, and vascular cells of the cerebral cortex. J Neurosci 34(36), 11929–11947. doi: 10.1523/JNEUROSCI.1860-14.2014.

